# ATG5 suppresses type I IFN-dependent neutrophil swarming and NETosis

**DOI:** 10.1101/2023.03.18.533244

**Authors:** Rachel L. Kinsella, Chanchal Sur Chowdhury, Asya Smirnov, Yassin Mreyoud, Jacqueline M. Kimmey, Ekaterina Esaulova, Darren Kreamalmeyer, Maxim N. Artyomov, Christina L. Stallings

## Abstract

Inflammation is critical for controlling infections, but when left unchecked can cause tissue damage and disease. For tuberculosis, the leading cause of death due to infection^1^, host inflammation is responsible for the clinical symptoms^2^, morbidity^2^, and mortality^3,4^. Specifically, neutrophil-dominated inflammation is associated with tuberculosis disease progression^3,5,6^. Therefore, understanding how neutrophil functions are regulated during infection is important for developing ways to prevent disease. *Atg5* was the first gene shown to specifically function within neutrophils to promote control of *Mycobacterium tuberculosis*^7^, the causative agent of tuberculosis. ATG5 is best studied for its role in autophagy^8–11^, however, the protective activity of ATG5 in neutrophils was unexpectedly independent of other autophagy proteins and remained elusive^7^. We report here that ATG5, but not other autophagy proteins, is required in neutrophils to suppress neutrophil NETosis and swarming that occur due to elevated type I interferon levels during *M. tuberculosis* infection. The elevated level of NETosis that results from loss of ATG5 expression contributes to the early susceptibility of *Atg5*^*fl/fl*^*-LysM-Cre* mice during *M. tuberculosis* infection. NETosis is associated with poor disease outcomes in tuberculosis^12,13^ and COVID-19 patients^14,15^, as well as during other inflammatory diseases in humans^16,17^. Our studies identify an essential regulator of NETosis and elucidate previously unappreciated roles for ATG5 during infection, which may inform the design of host-directed therapeutics modulating these pathways.

## MAIN TEXT

*Atg5*^*fl/fl*^*-LysM-Cre* mice, which delete the *Atg5* gene specifically in macrophages, inflammatory monocytes, some dendritic cells, and neutrophils, are severely susceptible to infection with a standard low-dose (∼100 colony forming units (CFU)) of aerosolized *M. tuberculosis* and succumb to infection by 40 days post-infection (dpi)^7,18,19^. *Atg5*^*fl/fl*^*-LysM-Cre* mice develop neutrophil dominated lung inflammation by 14 dpi that contributes to their susceptibility, where neutrophil depletion extends survival^7^. In addition, loss of ATG5 expression specifically in neutrophils is sufficient to cause susceptibility to *M. tuberculosis* infection^7^. The essential role for ATG5 in neutrophils to control *M. tuberculosis* pathogenesis is independent of other autophagy proteins, leaving open the question of how ATG5 functions in neutrophils to control *M. tuberculosis* infection. To determine how loss of ATG5 affects neutrophil responses during *M. tuberculosis* infection, we performed RNA sequencing (RNA-seq) analysis of neutrophils isolated from the lungs of *Atg5*^*fl/fl*^*-LysM-Cre* and *Atg5*^*fl/fl*^ mice at 14 dpi **(Table S1)** and identified the pathways enriched for genes differentially expressed in the absence of *Atg5* by performing Gene Set Enrichment Analysis (GSEA) on the genes ranked as a function of p value and log fold change (qnorm(1-PValue/2)*sign(logFC) in *Atg5*^*fl/fl*^*-LysM-Cre* versus *Atg5*^fl/fl^ neutrophils using the HALLMARK gene sets from the Molecular Signatures Database^20,21^ and the Reactome gene sets from the Reactome.org project^22^. GSEA revealed that pathways that were under-enriched for expression in neutrophils from *M. tuberculosis* infected *Atg5*^*fl/fl*^*-LysM-Cre* mice compared to controls related to metabolism, including amino acid metabolism **(Fig. 1A)**. This is consistent with the canonical role of autophagy in amino acid metabolism and recycling^23–27^, where ATG5-deficient neutrophils would be defective in these pathways. In contrast, three of the ten most enriched pathways expressed in neutrophils from *M. tuberculosis*-infected *Atg5*^*fl/fl*^*-LysM-Cre* mice were related to type I interferon (IFN) and RIG-I **(Fig. 1A, B)**. These data suggest that loss of ATG5 in LysM^+^ cells results in amplified type I IFN signaling in neutrophils during *M. tuberculosis* infection. Type I IFN has been associated with poor control of *M. tuberculosis* infection in humans, where leukocytes from active TB patients express higher levels of type I IFN-inducible genes than latently infected or uninfected people^28,29^. Most reports involving *M. tuberculosis* infection of mice also support a role for type I IFN signaling in exacerbating *M. tuberculosis* infection^12,28,30–32^, although the mechanistic basis for how type I signaling interferes with control of *M. tuberculosis* remains unknown. To determine if the higher levels of type I IFN signaling in neutrophils from *M. tuberculosis*-infected *Atg5*^*fl/fl*^*-LysM-Cre* mice contributed to their susceptibility, we generated *Atg5*^*fl/fl*^*-LysM-Cre* and *Atg5*^*fl/fl*^ mice deleted for the gene encoding the type I interferon receptor, *Ifnar1*, and monitored survival following infection with an *M. tuberculosis* strain constitutively expressing GFP (GFP-*Mtb*^33,34^) **(Fig. 1C)**. Deletion of *Ifnar1* significantly extended the median survival time of *Atg5*^*fl/fl*^*-LysM-Cre* mice from 30 dpi to 136 dpi, indicating that increased type I IFN signaling contributes to the susceptibility of *Atg5*^*fl/fl*^*-LysM-Cre* mice to *M. tuberculosis*. The *Ifnar1*^-/-^/*Atg5*^*fl/fl*^*-LysM-Cre* mice still succumbed to *M. tuberculosis* infection significantly earlier than *Atg5*^*fl/fl*^ (median survival 253 dpi) and *Ifnar1*^*-/-*^*/Atg5*^*fl/fl*^ (median survival 271 dpi) mice, indicating that there are also type I IFN-independent effects of loss of ATG5 impacting survival **(Fig 1C)**.

**Figure 1.**
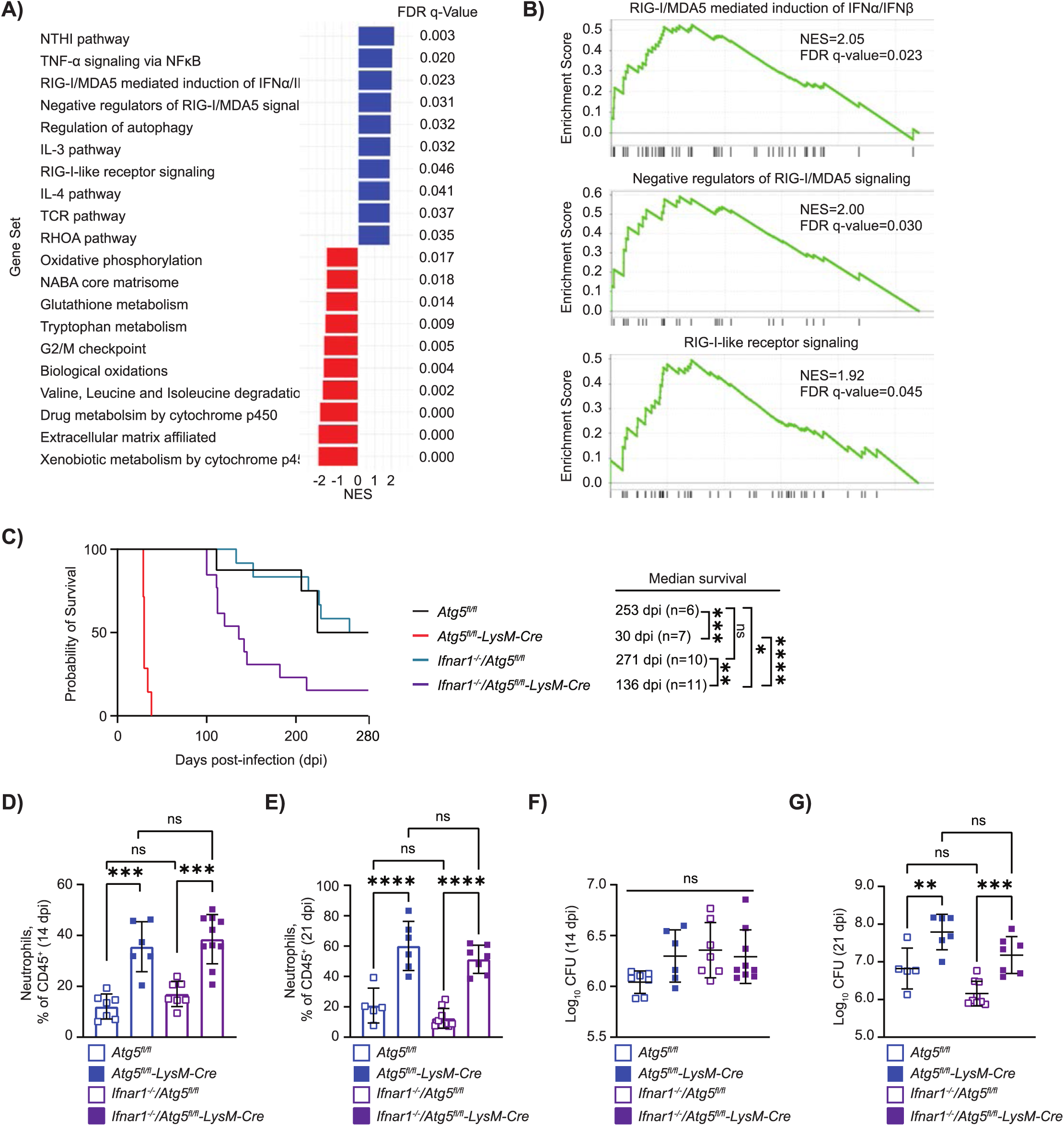
Loss of type I IFN signaling rescues the susceptibility of *Atg5*^fl/fl^-*LysM-Cre* mice to *M. tuberculosis* infection without affecting bacterial burdens or neutrophil accumulation. **(A)** Gene set enrichment analysis (GSEA) of transcripts from neutrophils isolated at 14 dpi from the lungs of *M. tuberculosis* infected mice showing the normalized enrichment score (NES) for the top ten pathways enriched (blue) or under-enriched (red) in *Atg5*^*fl/fl*^*-LysM-Cre* neutrophils compared to *Atg5*^*fl/fl*^ neutrophils. Differentially expressed genes were identified using a Benjamini-Hochberg adjustment with an adjusted p-value < 0.05 and expressed as an FDR q-value. **(B)** GSEA enrichment plots of type I interferon and RIG-I related pathways in neutrophils from *Atg5*^*fl/fl*^*-LysM-Cre* mice compared to *Atg5*^*fl/*fl^ at 14 dpi. **(C)** Kaplan Meier curve of survival proportions during *M. tuberculosis* infection of mice. The median survival time for each genotype is reported. **(D-E)** Proportion of CD45^+^ cells that are neutrophils (Ly6G^+^ CD11b^+^) in the lungs of mice at **(D)** 14 dpi and **(E)** 21 dpi. **(F-G)** Log transformed graph of the colony forming units (CFU) in the right lung of mice at **(F)** 14 and **(G)** 21 dpi. **(D-G)** Bar graph of data represents mean ± SD and each data point is from an individual mouse compiled from at least two separate infection experiments. Statistical differences were determined by a log-rank Mantel-Cox test **(C)** or one-way ANOVA and Šídák multiple comparison test **(D-G)**. * P < 0.05, ** P < 0.01, *** P < 0.001, **** P < 0.0001. Differences that are not statistically significant are designated as ns.

We evaluated lung bacterial burden and neutrophil accumulation at 14 and 21 dpi in *Ifnar1*^-/-^/*Atg5*^*fl/fl*^*- LysM-Cre, Atg5*^*fl/fl*^*-LysM-Cre, Ifnar1*^*-/-*^*/Atg5*^*fl/fl*^, and *Atg5*^*fl/fl*^ mice to determine whether the improved survival in the absence of type I IFN signaling was associated with effects on pathogen replication and inflammation. *Ifnar1*^*-/-*^/*Atg5*^*fl/fl*^*-LysM-Cre* mice accumulated a similarly high frequency of neutrophils in the lungs at 14 and 21 dpi as *Atg5*^*fl/fl*^*-LysM-Cre* mice **(Fig. 1D, E)**. In addition, *M. tuberculosis* burden was similarly high in *Ifnar1*^*-/-*^/*Atg5*^*fl/fl*^*-LysM-Cre* mice and *Atg5*^*fl/fl*^*-LysM-Cre* mice at 21 dpi, demonstrating that type I IFN signaling was not directly affecting *M. tuberculosis* replication **(Fig. 1F, G)**. The elevated *M. tuberculosis* burden observed in *Ifnar1*^*-/-*^/*Atg5*^*fl/fl*^*-LysM-Cre* mice, which still accumulate high numbers of neutrophils, is consistent with our previous findings that the enhanced recruitment of neutrophils in *Atg5*^*fl/fl*^*-LysM-Cre* mice promotes *M. tuberculosis* growth^7^. However, we have also previously shown that neutrophilia directly contributes to the susceptibility of *Atg5*^*fl/fl*^*-LysM-Cre* mice to *M. tuberculosis* infection, as depletion of neutrophils significantly prolongs survival in these mice^7^.

Therefore, the finding that deletion of *Ifnar1* rescues the susceptibility of *Atg5*^*fl/fl*^*-LysM-Cre* mice without affecting neutrophil recruitment implies that type I IFN regulates neutrophil activity during *M. tuberculosis* infection.

Type I IFN signaling in neutrophils has been linked to the production of neutrophil extracellular traps (NETs)^12,16,35–38^. NETosis is the process whereby neutrophils undergo histone citrullination, which leads to the chromatin decondensation that is required for the release of web-like chromatin structures decorated with antimicrobial granule proteins with the potential to bind, kill, and trap pathogens^39^. Despite their canonical antimicrobial role, NETs do not kill *M. tuberculosis* and there is evidence that NETs promote *M. tuberculosis* pathogenesis^40,41^. NETosis has also recently been associated with necrotic tuberculosis lesions showing poor responses to antibiotic therapy in humans^12^. To determine if the higher levels of type I IFN signaling was accompanied by increased NETosis in *M. tuberculosis* infected *Atg5*^*fl/fl*^*-LysM-Cre* mice, we analyzed lung lesions of *Atg5*^*fl/fl*^*-LysM-Cre* and *Atg5*^*fl/fl*^ mice at 14 and 21 dpi for structural features consistent with NETs using scanning electron microscopy (SEM) **(Fig. 2A)**. At 14 dpi, we observed NET-like structures in the lung lesions of *Atg5*^*fl/fl*^*-LysM-Cre* mice that were not as apparent in the *Atg5*^*fl/fl*^ mice. At 21 dpi, the NET-like strands were visually more abundant in the lesions of *Atg5*^*fl/fl*^*-LysM-Cre* than *Atg5*^*fl/fl*^ mice. To specifically determine if the NET-like structures observed by SEM represented higher levels of NETosis in *M. tuberculosis* infected *Atg5*^*fl/fl*^*-LysM-Cre* mice, we performed immunofluorescence histology of lung tissue sections at 21 dpi **(Fig. 2B)**. Neutrophils were identified by staining with anti-Ly6G antibody and NETosis was monitored using an antibody specific for citrullination of histone 3 (H3Cit), a marker of the initial step of NETosis^12,42,43^. Compared to *Atg5*^*fl/fl*^ mice, the percentage of H3Cit^+^ area per lung section was significantly higher in *Atg5*^*fl/fl*^*-LysM-Cre* mice at 21 dpi **(Fig. 2C)**, indicating that there are more neutrophils undergoing NETosis. Since *Atg5*^*fl/fl*^*-LysM-Cre* mice have a higher number of neutrophils in the lung at 21 dpi compared to *Atg5*^*fl/fl*^ mice **(Fig. 1E)**^7^, we also calculated the percentage Ly6G^+^ area that was H3Cit^+^ to standardize for the number of neutrophils **(Fig. 2D)**. The percent of total Ly6G^+^ area that was H3Cit^+^ in *Atg5*^*fl/fl*^*-LysM-Cre* lung sections was significantly higher than in *Atg5*^*fl/fl*^ sections at 21 dpi **(Fig. 2D)**, indicating that ATG5-deficient neutrophils were more likely to be undergoing NETosis. We also measured neutrophil histone citrullination in the lungs of *Atg5*^*fl/fl*^*-LysM-Cre* and *Atg5*^*fl/fl*^ mice at 21 dpi by flow cytometry and observed a significantly higher number of H3Cit^+^ neutrophils and higher proportion of total neutrophils that were H3Cit^+^ in the lungs of *Atg5*^*fl/fl*^*-LysM-Cre* mice **(Fig. 2E-G)**, further demonstrating that loss of ATG5 in LysM^+^ cells results in a greater propensity for neutrophils to undergo NETosis during *M. tuberculosis* infection *in vivo*.

**Figure 2.**
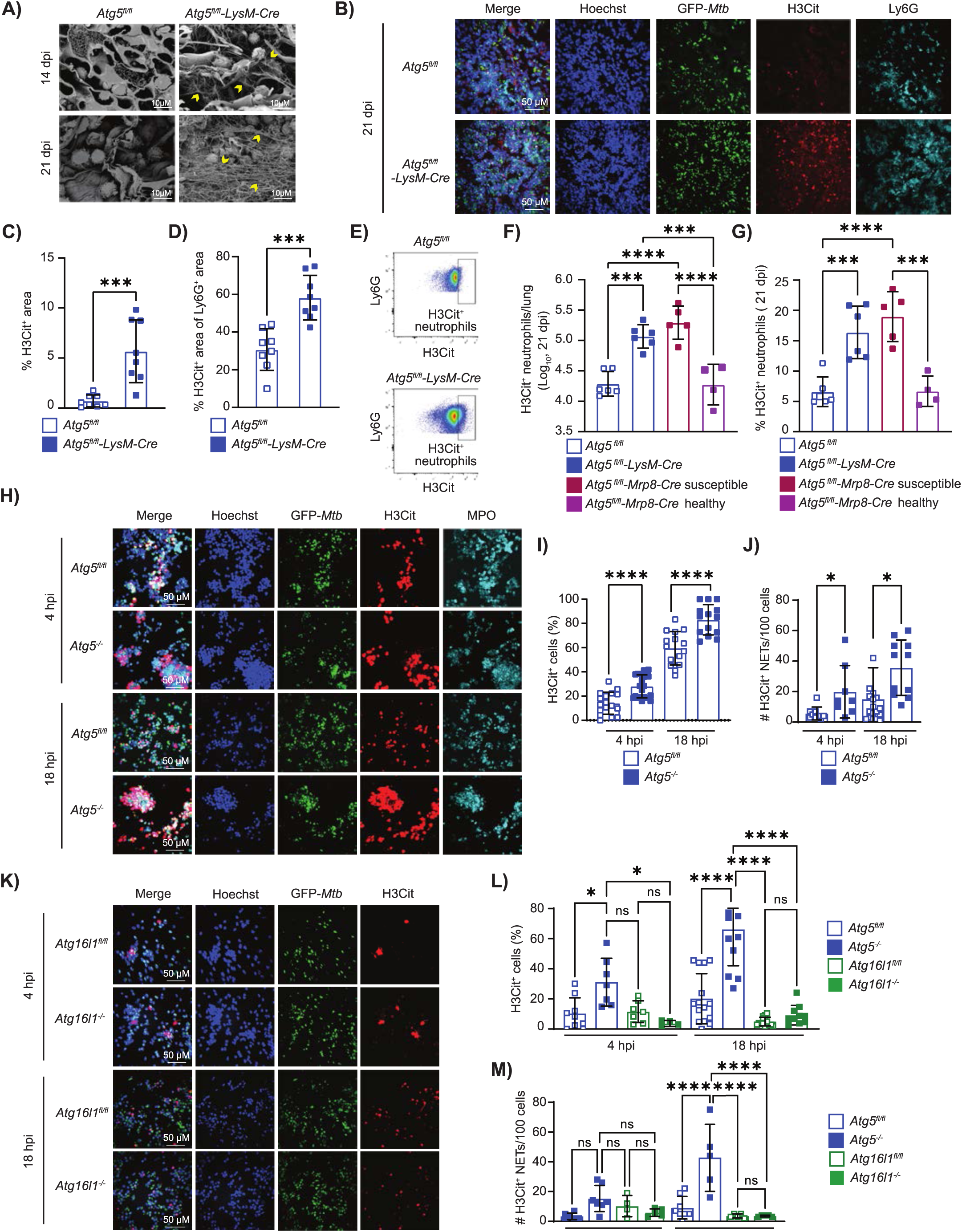
ATG5 functions in neutrophils independent of autophagy to suppress NETosis *in vivo* and *in vitro*. **(A)** Representative scanning electron microscopy images of GFP-*Mtb* infected mouse lungs at 14 and 21 dpi. Neutrophil extracellular trap (NET)-like structures are indicated by yellow arrows. Images were acquired at 2000X magnification. **(B)** Representative confocal immunofluorescence images of lung sections from GFP-*Mtb* infected mice at 21 dpi that were probed with antibodies to detect citrullinated histone H3 (H3Cit; red), Ly6G (neutrophil marker; cyan), and DNA (Hoechst; blue). GFP-*Mtb* is also shown. **(C)** The % of H3Cit^+^ pixel area per field and **(D)** the percent of Ly6G^+^ pixel area that was H3Cit^+^ per lung section in confocal immunofluorescence images of lung sections from GFP-*Mtb* infected mice at 21 dpi. **(C**,**D)** Each data point is from one field of view and two fields of view per lung histology section from 4 mice from at least two separate infection experiments were compiled. **(E)** Representative flow cytometry plot of H3Cit^+^ neutrophils (CD45^+^Ly6G^+^CD11b^+^) from the lungs of mice at 21 dpi. **(F)** The number of H3Cit^+^ neutrophils and **(G)** the frequency of neutrophils that are H3Cit^+^ at 21 dpi in the lungs of mice. *Atg5*^*fl/fl*^*-Mrp8-Cre* mice were categorized as susceptible if they have lost >5% of their weight by 20 dpi and healthy if they have not. **(F-G)** Each data point is from an individual mouse compiled from at least two separate infection experiments. **(H**,**K)** Representative confocal immunofluorescence microscopy images of bone marrow neutrophils infected with GFP-*Mtb in vitro* for 4 or 18 hours and stained for H3Cit, the neutrophil marker myeloperoxidase (MPO), and DNA (Hoechst) (60X magnification). **(I**,**L)** The proportion of Hoechst^+^ cells that were H3Cit^+^ and **(J**,**M)** the number of extracellular H3Cit^+^ NETs released per 100 cells in fluorescent microscopy images of neutrophils at 4 and 18 hpi with GFP-*Mtb*. **(I-J, L-M)** Each datapoint represents one field and a minimum of 6 fields containing 40-150 cells/field compiled from at least 2 independent experiments has been graphed. All graphs report the mean ± SD. Statistical differences were determined by student t-test **(C-D, I-J** comparing only within a single timepoint) or one-way ANOVA and Šídák multiple comparison test **(F-G, L-M)**. * P < 0.05, *** P < 0.001, **** P < 0.0001. Differences that are not statistically significant are designated as ns.

To determine if the role for ATG5 in suppressing NETosis *in vivo* was neutrophil intrinsic, we analyzed H3Cit positivity in *M. tuberculosis* infected *Atg5*^*fl/fl*^*-Mrp8-Cre* mice, which delete *Atg5* specifically in neutrophils. We previously showed that loss of *Atg5* in neutrophils results in increased susceptibility to *M. tuberculosis* infection in some, but not all, *Atg5*^*fl/fl*^*-Mrp8-Cre* mice^7^. If we separate the susceptible and healthy *Atg5*^*fl/fl*^*-Mrp8-Cre* mice as done previously where susceptible *Atg5*^*fl/fl*^*-Mrp8-Cre* mice have lost more than 5% of their pre-infection body weight by 20 dpi^7^, the number of H3Cit^+^ neutrophils and the proportion of total neutrophils that are H3Cit^+^ was significantly higher in susceptible *Atg5*^*fl/fl*^*-Mrp8-Cre* mice relative to *Atg5*^*fl/fl*^ mice at 21 dpi **(Fig. 2F**,**G)**. These data indicate that loss of *Atg5* specifically in neutrophils can result in higher levels of NETosis, implying there is a neutrophil intrinsic role for ATG5 in suppressing NETosis during *M. tuberculosis* infection *in vivo*. To probe this neutrophil intrinsic role for ATG5 further, we isolated neutrophils from the bone marrow of *Atg5*^*fl/fl*^*-LysM-Cre* and *Atg5*^*fl/fl*^ mice, infected the neutrophils with GFP-*Mtb in vitro*, and monitored NETosis by immunofluorescence microscopy **(Fig. 2H)**. By 4 hours post infection (hpi), we observed a significantly higher frequency of H3Cit^+^ neutrophils in *Atg5*^*-/-*^ cultures compared to *Atg5*^*fl/fl*^ cultures **(Fig. 2I)**. The higher frequency of H3Cit^+^ neutrophils in *Atg5*^*-/-*^ cultures was maintained at 18 hpi **(Fig. 2I)**. In addition to monitoring histone citrullination, we also quantified the number of released NETs by counting extracellular H3Cit^+^ web-like structures using the Ridge Detection plugin^44^ for the Fiji image analysis program^45^ **(Fig. S1A)**. At both 4 and 18 hpi, *Atg5*^*-/-*^ neutrophils were associated with a greater number of released NETs than *Atg5*^*fl/fl*^ neutrophils **(Fig. 2J)**. We confirmed that the H3Cit^+^ webs represented released NETs by performing SEM of infected neutrophil cultures, where we observed significantly more NETs being released in *Atg5*^*-/-*^ neutrophil cultures compared to *Atg5*^*fl/fl*^ neutrophils at 4 and 18 hpi **(Fig. S1B-C)**. There was no difference in *M. tuberculosis* burdens in the *Atg5*^*-/-*^ and *Atg5*^*fl/fl*^ cultures at 4 or 18 hpi, where both time points are shorter than the doubling time for *M. tuberculosis*, demonstrating that the increased NETosis was not a result of more antigen being present **(Fig. S1D)**. Mock-infection did not cause *Atg5*^*-/-*^ neutrophils to NETose more than *Atg5*^*fl/fl*^ neutrophils, where on average less than 2% of neutrophils were H3Cit^+^ and released fewer than one H3Cit^+^ NET per 100 cells in cultures from both genotypes at 18 hours **(Fig. S1E-G)**. Together, these data support a neutrophil-intrinsic role for ATG5 in suppressing histone citrullination and NET release during *M. tuberculosis* infection. Infection of neutrophils lacking BECLIN 1 or ATG16L1, two additional proteins required for autophagy, with *M. tuberculosis* did not result in increased NETosis as compared to controls **(Fig. 2K-M and Fig. S1H-J)**, suggesting that this role for ATG5 in suppressing NETosis during *M. tuberculosis* infection is autophagy-independent. There was no difference in *M. tuberculosis* burdens in the *Atg5*^*-/-*^, *Becn1*^*-/-*^ and *Atg16l1*^*-/-*^ cultures at 4 or 18 hpi, demonstrating that increased NETosis in *Atg5*^*-/-*^ neutrophils was not a result of increased infection **(Fig. S1D)**.

Histone citrullination by protein arginine deiminase (PAD) enzymes is critical for the initial chromatin decondensation that is required for chromatin to be packaged into vesicles and released as NETs following most stimuli^39,46,47^, including during *M. tuberculosis* infection^40,48^. PAD4 is the primary PAD enzyme expressed in neutrophils and has been shown to be essential for NETosis in response to a number of different stimuli^39,46,47,49^. To determine whether PAD4 was responsible for the increased histone citrullination in *Atg5*^*-/-*^ neutrophils during *M. tuberculosis* infection, we crossed mice deleted for the gene encoding PAD4 (*Padi4*) with *Atg5*^*fl/fl*^*-LysM-Cre* and *Atg5*^*fl/fl*^ mice. Histone citrullination and released extracellular H3Cit^+^ NETs were almost undetectable in cultures of neutrophils isolated from *Padi4*^-/-^/*Atg5*^*fl/fl*^ and *Padi4*^-/-^/*Atg5*^*fl/fl*^*-LysM-Cre* mice following infecting with *M. tuberculosis in vitro* **(Fig. 3A-C)**. Therefore, *M. tuberculosis-*induced histone citrullination and NETosis in *Atg5*^*fl/fl*^ and *Atg5*^*-/-*^ neutrophils is *Padi4-*dependent. Deletion of *Padi4* in *Atg5*^*-/-*^ neutrophils had no impact on *M. tuberculosis* burden at 4 and 18 hpi, demonstrating that NETosis does not impact bacterial survival in these conditions up to 18 hpi **(Fig. S2A)**.

**Figure 3.**
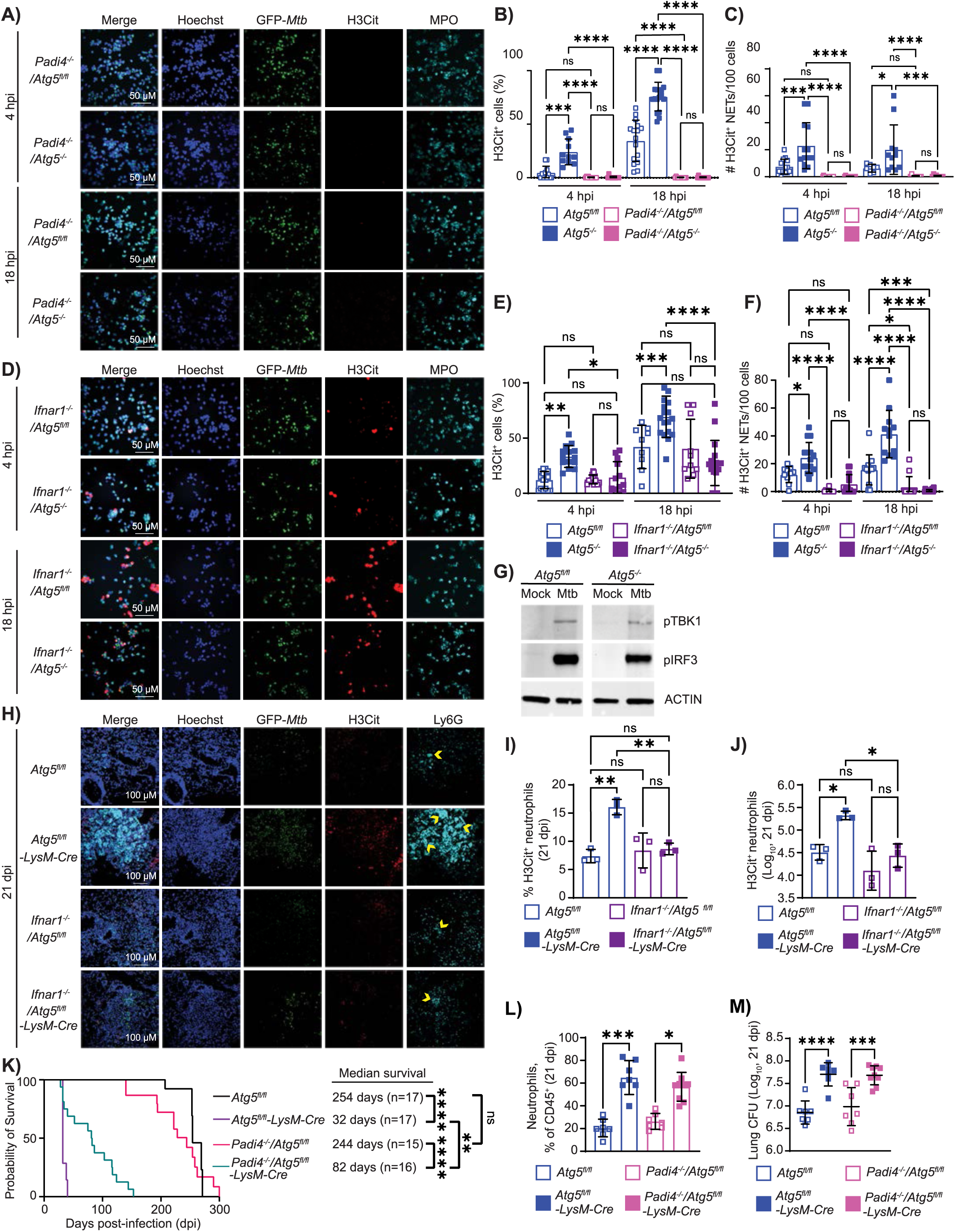
ATG5 suppresses type I IFN-induced PAD4-mediated histone citrullination during *M. tuberculosis* infection. (**A**,**D)** Representative confocal immunofluorescence images of mouse neutrophils infected with GFP-*Mtb in vitro* for 4 or 18 hpi and stained for H3Cit, neutrophil marker MPO, and DNA (Hoechst) (60X magnification). **(B**,**E)** The proportion of Hoechst^+^ cells that were H3Cit^+^ and **(C**,**F)** the number of extracellular H3Cit^+^ NETs released per 100 cells in fluorescent microscopy experiments. Bar graphs report the mean ± SD and each datapoint represents one field and a minimum of 6 fields containing 40-250 cells/field compiled from at least 2 independent experiments is graphed. **(G)** Western blots probing for phosphorylated TANK binding kinase 1 (pTBK1), phosphorylated interferon regulatory factor 3 (pIRF3) and ACTIN in *Atg5*^*fl/fl*^ and *Atg5*^*-/-*^ neutrophils mock treated or infected with GFP-*Mtb in vitro* at 4 hpi. **(H)** Representative confocal immunofluorescence images at 20X magnification of lung sections from GFP-*Mtb* infected mice at 21 dpi probed with antibodies to detect citrullinated histone H3 (H3Cit; red), neutrophil marker Ly6G (cyan), and DNA (Hoechst; blue). GFP-*Mtb* is also shown. Yellow arrows indicate Ly6G^+^ aggregates that were greater than 70 pixels in size. **(I)** The proportion of neutrophils (CD45^+^ Ly6G^+^ CD11b^+^) that were H3Cit^+^ and **(J)** the number of H3Cit^+^ neutrophils in the lungs of mice at 21 dpi measured by flow cytometry. The bar graphs of data represent mean ± SD, and each data point is from an individual mouse compiled from at least two separate infection experiments. **(K)** Kaplan Meier curve of survival proportions during GFP-*Mtb* infection of mice. The median survival time for each genotype is reported. **(L)** Proportion of CD45^+^ cells that are neutrophils (Ly6G^+^ CD11b^+^) in the lung at 21 dpi. **(M)** Log transformed graph of the CFU in the right lung of mice at 21 dpi. **(L-M)** Each data point is from an individual mouse compiled from at least two separate infection experiments and the mean ± SD is shown. Statistical differences were determined by one-way ANOVA and Šídák multiple comparison test except in **(K)** when a log-rank Mantel-Cox test was used. * P < 0.05, ** P < 0.01, *** P < 0.001, **** P < 0.0001. Differences that are not statistically significant are designated as ns.

To investigate whether type I IFN signaling directly contributed to the increased levels of histone citrullination and NET release in *M. tuberculosis* infected ATG5-deficient neutrophils, we isolated bone marrow neutrophils from *Ifnar1*^*-/-*^*/Atg5*^*fl/fl*^*-LysM-Cre, Ifnar1*^*-/-*^*/Atg5*^*fl/fl*^, *Atg5*^*fl/fl*^*-LysM-Cre*, and *Atg5*^*fl/fl*^ mice, infected with GFP-*Mtb* for 4 and 18 hours, and monitored NETosis by immunofluorescence microscopy **(Fig. 3D)**. In WT neutrophils, type I IFN signaling has no effect on PAD4-mediated histone citrullination during *M. tuberculosis* infection but instead promotes the packaging of decondensed chromatin into vesicles and subsequent NET release following histone citrullination^12,40^. Therefore, as expected, there was no significant difference in the levels of histone citrullination in *Atg5*^*fl/fl*^ and *Ifnar1*^*-/-*^*/Atg5*^*fl/fl*^ neutrophils, but a significantly decreased number of extracellular released NETs from *Ifnar1*^*-/-*^*/Atg5*^*fl/fl*^ neutrophils compared to *Atg5*^*fl/fl*^ neutrophils at 4 and 18 hpi **(Fig. 3E-F)**. In contrast, the increased H3Cit signal in *M. tuberculosis* infected *Atg5*^*-/-*^ neutrophils was reversed back to levels observed in *Atg5*^fl/fl^ neutrophils when *Ifnar1* was deleted **(Fig. 3E)**. Loss of type I IFN signaling in *Atg5*^*-/-*^ neutrophils also resulted in a decrease in H3Cit^+^ NETs released to levels observed in *Atg5*^*fl/fl*^ neutrophils **(Fig. 3F)**. There were no differences in *M. tuberculosis* CFU in *Ifnar1*^*-/-*^*/Atg5*^*-/-*^, *Ifnar1*^*-/-*^*/Atg5*^*fl/fl*^, *Atg5*^*-/-*^ and *Atg5*^*fl/fl*^ neutrophil cultures at 18 hpi **(Fig. S2B)**, therefore, the effects on NETosis are not a result of differences in *M. tuberculosis* burden. These data indicate that ATG5 specifically blocks type I IFN-induced PAD4 activity, where the ability of type I IFN signaling to promote PAD4 activity in neutrophils could only be observed when *Atg5* was deleted. To investigate whether ATG5 blocks type I IFN activation of PAD4 activity by suppressing the production of type I IFN during *M. tuberculosis* infection, we monitored the levels of phosphorylated TANK binding kinase 1 (pTBK1) and phosphorylated interferon regulatory factor 3 (pIRF3) in *Atg5*^*-/-*^ and *Atg5*^*fl/fl*^ neutrophil cultures at 4 hpi **(Fig. 3G)**. TBK1 and IRF3 are phosphorylated to activate transcription of type I IFNs in response to infection. We observed similar levels of pTBK1 and pIRF3 in *Atg5*^*-/-*^ and *Atg5*^*fl/fl*^ neutrophils at 4 hpi, suggesting that ATG5 functions downstream of type I IFN production to block type I IFN-induced PAD4 activity in neutrophils following *M. tuberculosis* infection. We directly tested the specificity of ATG5’s role in suppressing NETosis downstream of type I IFN by treating *Ifnar1*^*-/-*^*/Atg5*^*-/-*^, *Ifnar1*^*-/-*^*/Atg5*^*fl/fl*^, *Atg5*^*-/-*^, and *Atg5*^*fl/fl*^ neutrophils with ionomycin, an ionophore that induces histone citrullination and NETosis independent of type I IFN^50,51^. Neither deletion of *Ifnar1* nor *Atg5* affected the levels of histone citrullination or the number of released NETs following ionomycin treatment **(Fig. S2C-E)**, confirming that ATG5 specifically suppresses type I IFN-induced PAD4 activity. To determine if type I IFN signaling was responsible for the higher levels of histone citrullination in the neutrophils of *M. tuberculosis* infected *Atg5*^*fl/fl*^*-LysM-Cre* mice, we performed immunofluorescence histology and flow cytometry analysis to detect the levels of H3Cit^+^ cells in the lungs from *Atg5*^*fl/fl*^, *Atg5*^*fl/fl*^*-LysM-Cre, Ifnar1*^*-/-*^*/Atg5*^*fl/fl*^, and *Ifnar1*^*-/-*^*/Atg5*^*fl/fl*^*-LysM-Cre* mice at 21 dpi with GFP-*Mtb*. Deletion of *Ifnar1* reduced the higher levels of histone citrullination in the lung tissue sections of *Atg5*^*fl/fl*^*-LysM-Cre* mice at 21 dpi **(Fig. 3H)** and quantification by flow cytometry confirmed that there was a significantly decreased number and frequency of H3Cit^+^ neutrophils in the lungs of *Ifnar1*^*-/-*^*/Atg5*^*fl/fl*^*-LysM-Cre* mice compared to *Atg5*^*fl/fl*^*-LysM-Cre* mice at 21 dpi **(Fig. 3I-J)**. Therefore, ATG5 suppresses type I IFN-induced histone citrullination in neutrophils during *M. tuberculosis* infection *in vitro* and *in vivo*.

To determine if increased NETosis in *M. tuberculosis* infected *Atg5*^*fl/fl*^*-LysM-Cre* mice contributed to their susceptibility, we infected *Atg5*^*fl/fl*^, *Atg5*^*fl/fl*^*-LysM-Cre, Padi4*^*-/-*^*/Atg5*^*fl/fl*^, and *Padi4*^*-/-*^*/Atg5*^*fl/fl*^*-LysM-Cre* mice with GFP-*Mtb* and monitored survival. Deletion of *Padi4* significantly extended the median survival time of *Atg5*^*fl/fl*^*-LysM-Cre* mice from 32 dpi to 82 dpi during *M. tuberculosis* infection **(Fig. 3K)**. We confirmed that deletion of *Padi4* reduced histone citrullination in the lung during *M. tuberculosis* infection at 21 dpi by immunofluorescence histology and flow cytometry **(Fig. S2F-G)**. However, similar to deletion of *Ifnar1*^*-/-*^, *Padi4* deletion did not reduce neutrophil abundance or lung bacterial burden at 21dpi in *Atg5*^*fl/fl*^*-LysM-Cre* mice **(Fig. 3L-M)**. Therefore, the improved survival was not due to reduced neutrophil accumulation, but instead due to the decreased neutrophil NETosis during *M. tuberculosis* infection. These data indicate that increased NETosis is one way type I IFN confers susceptibility in *M. tuberculosis* infected *Atg5*^*fl/fl*^*-LysM-Cre* mice. However, the *Ifnar1*^*-/-*^*/Atg5*^*fl/fl*^*-LysM-Cre* mice (median survival time 136 dpi **(Fig. 1C)**) survived significantly (p=0.0027) longer than the *Padi4*^*-/-*^*/Atg5*^*fl/fl*^*-LysM-Cre* mice (median survival time 82 dpi **(Fig. 3K)**) following *M. tuberculosis* infection, indicating that NETosis is not the only way type I IFN signaling contributes to the susceptibility of *M. tuberculosis* infected *Atg5*^*fl/fl*^*-LysM-Cre* mice. During the immunofluorescence microscopy analysis of *Atg5*^*fl/fl*^ and *Atg5*^*-/-*^ neutrophils infected with *M. tuberculosis in vitro*, we noted that *Atg5*^*-/-*^ neutrophils aggregated into significantly larger clusters than *Atg5*^*fl/fl*^ neutrophils by 4 hpi **(Fig. 2H)**, which we quantified using Fiji Particle Detection analysis of Hoechst^+^ cells **(Fig. 4A, Fig. S3A)**. The three-dimensional nature of the neutrophil aggregates in *M. tuberculosis* infected *Atg5*^*-/-*^ neutrophil cultures was even more apparent by SEM **(Fig. 4B)**. Mock infection of *Atg5*^*-/-*^ neutrophils did not result in significantly larger aggregates than mock-infected *Atg5*^*fl/fl*^ neutrophils **(Fig. S3B)**, demonstrating that the hyper-aggregation in the absence of ATG5 is dependent on infection. In contrast to ATG5^-/-^ neutrophils, *M. tuberculosis* infection of *Becn1*^*-/-*^ and *Atg16l1*^*-/-*^ neutrophils did not result in larger aggregates than *Becn1*^*fl/fl*^ and *Atg16l1*^*fl/fl*^ control neutrophils at 4hpi **(Fig. 4C)**. Therefore, in addition to suppressing NETosis, ATG5 functions independent of autophagy in neutrophils to inhibit aggregation during *M. tuberculosis* infection.

**Figure 4.**
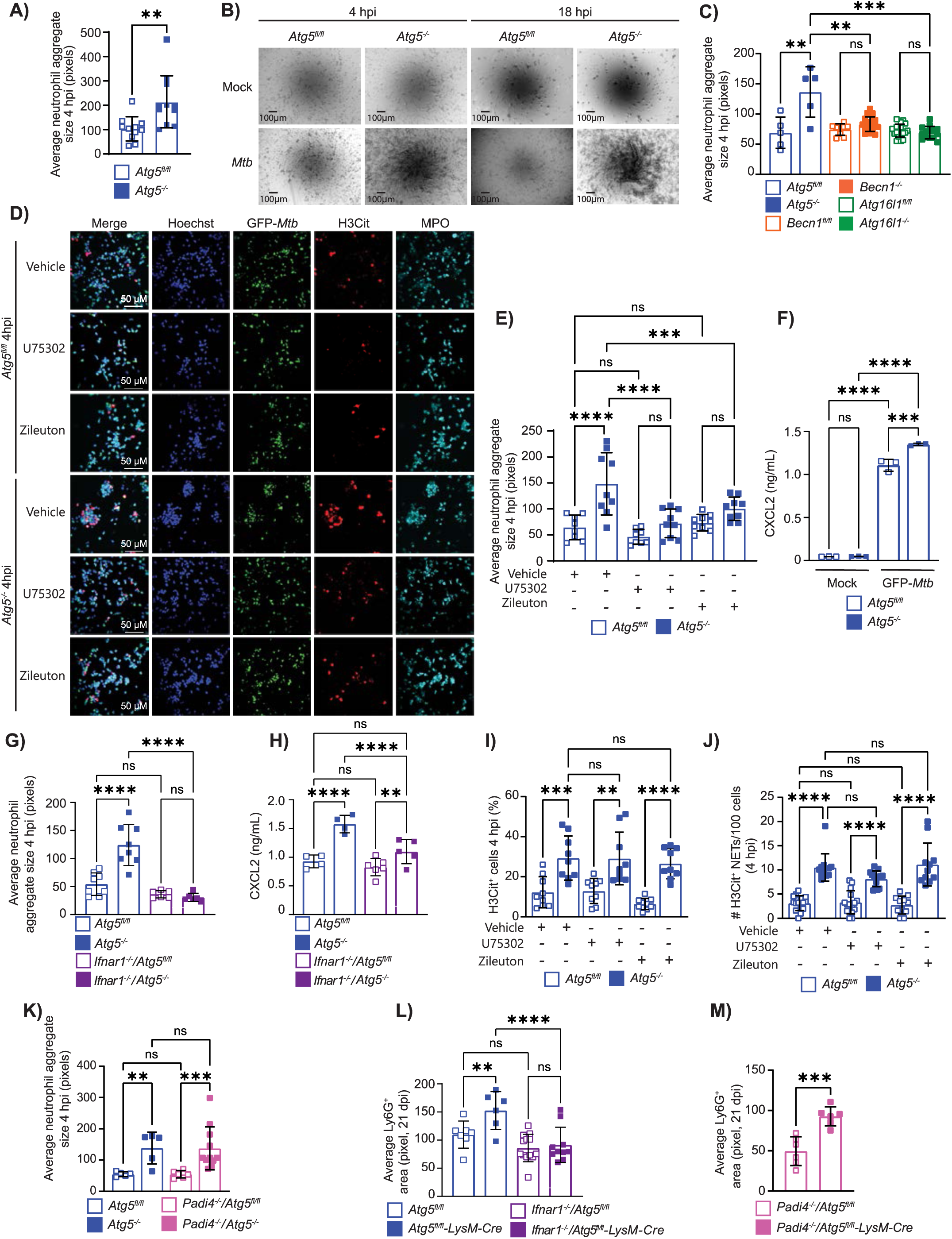
ATG5 suppresses type I IFN-induced neutrophil swarming independently of the regulation of NETosis. (**A**,**C**,**E**,**G**,**K)** Neutrophil aggregate sizes in fluorescence microscopy of mouse neutrophils infected with GFP-*Mtb in vitro* for 4 hpi. Each datapoint represents the average neutrophil aggregate size per field and a minimum of 5 fields containing 40-250 cells/field compiled from at least 2 independent experiments is graphed. **(B)** Representative SEM images at 100X magnification of bone marrow neutrophils mock or GFP-*Mtb* infected for 4 and 18 hours. **(D)** Representative confocal immunofluorescence microscopy images of mouse neutrophils treated with vehicle, U75302, or Zileuton and infected with GFP-*Mtb in vitro* for 4 hpi. Cells were stained for H3Cit, DNA (Hoechst) and neutrophil marker MPO (60X magnification). **(F**,**H)** The amount of CXCL2 present in neutrophil culture supernatants from mock treated or GFP-*Mtb* infected mouse neutrophils at 4 hpi quantified by ELISA. Each datapoint is from an independent well of infected cells compiled from at least 2 independent experiments. **(I)** The proportion of Hoechst^+^ cells that are H3Cit^+^ and **(J)** the number of extracellular H3Cit^+^ NETs released per 100 cells from neutrophils treated with vehicle, U75302 or Zileuton and infected with GFP-*Mtb in vitro* for 4 hpi, calculated from fluorescence microscopy experiments. **(L-M)** Average size of Ly6G^+^ areas within lung sections at 21 dpi (20x magnification) in GFP-*Mtb* infected mice. Each data point is the average pixel size of Ly6G^+^ aggregates in a single field, where lung sections from at least 6 mice compiled from at least two separate infection experiments were imaged. All graphs report the mean ± SD. Statistical differences were determined by student t-test **(A**,**M)** or one-way ANOVA and Šídák multiple comparison test **(C, E-L)**. ** P < 0.01, *** P < 0.001, **** P < 0.0001. Differences that are not statistically significant are designated as ns.

The aggregates of *Atg5*^*-/-*^ neutrophils are reminiscent of neutrophil swarms^52,53^. Neutrophil swarming is coordinated neutrophil migratory behavior in response to chemoattractant signaling from leukotriene B4 (LTB4)^54–56^, CXCL2^54,57^, and formyl peptides^54^ that amplifies the recruitment of patrolling neutrophils to sites of damage or infection^54,57–62^. Efficient termination of neutrophil swarming is necessary to protect against tissue damage^58,59,63^, facilitate phagocytosis^62^, and contain bacterial infection^62,63^. To determine if *Atg5*^-/-^ neutrophils aggregated during *M. tuberculosis* infection due to increased neutrophil swarming, we blocked LTB4 signaling by treating with the leukotriene inhibitor Zileuton^52,64^ (Cayman Chemicals) or a leukotriene receptor antagonist U75302^52,65^ (Cayman Chemicals) during GFP-*Mtb* infection of *Atg5*^*-/-*^ and *Atg5*^*fl/fl*^ neutrophils and monitored neutrophil aggregation by immunofluorescence microscopy at 4 hpi **(Fig. 4D)**. Treatment with U75302 or Zileuton significantly reduced the average size of *Atg5*^*-/-*^ neutrophil aggregates to a similar size as observed in *Atg5*^*fl/fl*^ cultures at 4hpi **(Fig. 4E)**. These data demonstrate that LTB4 signaling is required for the aggregation behavior of *Atg5*^*-/-*^ neutrophils during *M. tuberculosis* infection, indicating that the aggregates represent increased neutrophil swarming. To determine if *Atg5*^*-/-*^ neutrophils form larger swarms during *M. tuberculosis* infection because of increased LTB4 or CXCL2 production by *Atg5*^*-/-*^ neutrophils, we quantified the levels of LTB4 and CXCL2 by enzyme linked immunosorbent assay (ELISA) **(Fig. 4F)**.

*M. tuberculosis* infection of both *Atg5*^*-/-*^ and *Atg5*^*fl/f*^ neutrophils induced LTB4 and CXCL2 production by 4 hours in comparison to mock treated neutrophils. The levels of LTB4 secreted by *M. tuberculosis* infected *Atg5*^*-/-*^ and *Atg5*^*fl/f*^ neutrophils at 4 hpi was similar **(Fig. S3C)**, however, the amount of CXCL2 secreted from *Atg5*^*-/-*^ neutrophils during *M. tuberculosis* infection was significantly higher than *Atg5*^*fl/fl*^ neutrophils **(Fig. 4F)**, which could contribute to the increased swarming.

To determine if the increased swarming by *Atg5*^-/-^ neutrophils was also induced by type I IFN, we quantified the size of neutrophil aggregates in cultures of neutrophils isolated from the bone marrow of *Atg5*^*fl/fl*^, *Atg5*^*fl/fl*^*-LysM-Cre, Ifnar1*^*-/-*^*/Atg5*^*fl/fl*^, and *Ifnar1*^*-/-*^*/Atg5*^*fl/fl*^*-LysM-Cre* mice and infected with *M. tuberculosis* for 4 hours *in vitro*. Deletion of *Ifnar1* reversed the larger sized aggregates formed by *Atg5*^*-/-*^ neutrophils during *M. tuberculosis* infection, decreasing the size of the aggregates to the same size as observed in *Atg5*^*fl/fl*^ neutrophil cultures **(Figs. 3D**,**4G)**. Given the association of increased CXCL2 production by *Atg5*^*-/-*^ neutrophils with the swarming phenotype, we investigated the effect of type I IFN on CXCL2 production by performing ELISAs on *Atg5*^*fl/fl*^, *Atg5*^*-/-*^, *Ifnar1*^*-/-*^*/Atg5*^*fl/fl*^, and *Ifnar1*^*-/-*^*/Atg5*^*-/-*^ neutrophil cultures infected *in vitro* with *M. tuberculosis* for 4 hours. *Ifnar1*^*-/-*^*/Atg5*^*-/-*^ neutrophils produced significantly less CXCL2 during *M. tuberculosis* infection than *Atg5*^*-/-*^ neutrophils **(Fig. 4H)**, together indicating that type I IFN signaling in *Atg5*^*-/-*^ neutrophils promotes CXCL2 production and swarming. Similar to our observations with histone citrullination, there was no difference in aggregate size or CXCL2 production between *Ifnar1*^*-/-*^*/Atg5*^*fl/fl*^ and *Atg5*^*fl/fl*^ neutrophils during *M. tuberculosis* infection **(Figs. 4G-H)**, further supporting that ATG5 suppresses type I IFN induced CXCL2 production and swarming.

Neutrophil swarming and NETosis have been observed in similar contexts and proposed to be associated^53,66^, however, a direct link has yet to be determined. To investigate if increased rates of NETosis in *M. tuberculosis* infected *Atg5*^*-/-*^ neutrophils were a result of the increased swarming, we measured histone citrullination and NET release following U75302 or Zileuton treatment using fluorescent microscopy at 4 hpi. U75302 or Zileuton treatment had no effect on the increased frequency of histone citrullination or NET release by *Atg5*^*-/-*^ neutrophils compared to *Atg5*^*fl/fl*^ neutrophils at 4 hpi **(Fig. 4D, I-J)**, demonstrating that the increased swarming in *M. tuberculosis* infected *Atg5*^*-/-*^ neutrophils is not responsible for the increased NETosis. Similarly, deleting *Padi4* to inhibit NETosis had no effect on swarming, where the size of *Padi4*^*-/-*^*/Atg5*^*-/-*^ neutrophil aggregates were similar to *Atg5*^*-/-*^ aggregates and significantly larger than *Padi4*^-/-^/*Atg5*^*fl/fl*^ and *Atg5*^*fl/fl*^ neutrophil aggregates at 4 hpi **(Fig. 4K)**. Therefore, the increased swarming of *M. tuberculosis* infected *Atg5*^*-/-*^ neutrophils is not a result of increased NETosis. Together, these data demonstrate that type I IFN and ATG5 regulate swarming and NETosis as two independent processes.

Since NETosis and swarming are both independently induced by type I IFN in the absence of ATG5, it is possible that both processes combine to cause the extreme susceptibility of *Atg5*^*fl/fl*^*-LysM-Cre* mice to *M. tuberculosis* infection. We noted that despite *Atg5*^*fl/fl*^*-LysM-Cre* and *Ifnar1*^*-/-*^*/Atg5*^*fl/fl*^*-LysM-Cre* mice accumulating a similarly high number of neutrophils in the lung at 21 dpi **(Fig. 1E)**, the physical distribution of the Ly6G^+^ neutrophils within the lung histology sections appeared different, where *Atg5*^*fl/fl*^*- LysM-Cre* mice exhibited larger Ly6G^+^ aggregates compared to *Ifnar1*^*-/-*^*/Atg5*^*fl/fl*^*-LysM-Cre* mice **(Fig. 3H, yellow arrows)**. We quantified the area of contiguous Ly6G^+^ signal using Fiji particle analysis to determine the size of the neutrophil aggregates in each lung histology section. Indeed, the neutrophil aggregates in the lungs of *Atg5*^*fl/fl*^*-LysM-Cre* mice were significantly larger than those observed in *Atg5*^*fl/fl*^ and *Ifnar1*^*-/-*^*/Atg5*^*fl/fl*^*-LysM-Cre* mice **(Fig. 4L)**. We hypothesize that the Ly6G^+^ aggregates in the lung histology sections represent neutrophil swarms that are induced by type I IFN in the absence of ATG5. Similar to the *in vitro* studies, there was no difference in neutrophil aggregate size in the lungs of *Ifnar1*^*-/-*^*/Atg5*^*fl/fl*^ and *Atg5*^*fl/fl*^ mice during *M. tuberculosis* infection **(Fig. 4L)**, further supporting that ATG5 suppresses type I IFN-induced neutrophil swarming during *M. tuberculosis* infection *in vivo*. Deletion of *Padi4* did not affect the size of neutrophil aggregates in *Atg5*^*fl/fl*^*-LysM-Cre* lung sections at 21 dpi **(Figs. S2F, 4M)**, supporting the *in vitro* data that type I IFN-dependent neutrophil swarming occurs independent of NETosis during *M. tuberculosis* infection. Given that deletion of *Ifnar1* in *Atg5*^*fl/fl*^*- LysM-Cre* mice extended survival further than *Padi4* deletion **(Figs. 1C, 3K)**, increased neutrophil swarming could be the additional way type I IFN exacerbates control of *M. tuberculosis* pathogenesis in the absence of ATG5.

Together, our data support a model where ATG5 is required in neutrophils to suppress type I IFN-induced PAD4-mediated histone citrullination and CXCL2 production during *M. tuberculosis* infection, two potential outcomes of type I IFN signaling that could only be revealed in the absence of ATG5 **(Fig. S3D)**. In the absence of ATG5, increased PAD4-mediated histone citrullination leads to uncontrolled NETosis and increased CXCL2 production is associated with heightened and persistent neutrophil swarming. Neutrophil swarming occurs in response to infection or cell damage when neutrophil-derived LTB4 recruits neutrophils from long distances to form transient aggregates and CXCL2 or fMLP signaling within the cluster promotes prolonged aggregation^54,55,57^. LTB4 has been shown to promote lung damage, inflammation, and susceptibility during *M. tuberculosis* infection^70,71^. In particular, survival can be extended in mice that are hypersusceptible to *M. tuberculosis* infection due to elevated type I IFN signaling by inhibiting LTB4 synthesis with Zileuton and addition of PGE2^72^, supporting a link between type I IFN and LTB4 associated pathways, such as swarming.

Our *in vitro* studies demonstrate that neutrophils produce type I IFN during *M. tuberculosis* infection and that ATG5 functions in neutrophils downstream of type I IFN signaling to suppress NETosis and swarming. The increased type I IFN signaling detected in lung neutrophils during *M. tuberculosis* infection of *Atg5*^*fl/fl*^*-LysM-Cre* mice implies that there is a positive feedback loop occurring that induces increased type I IFN production in the absence of ATG5 *in vivo*. The neutrophil-intrinsic role for ATG5 would also be expected to regulate neutrophil responses to type I IFN produced by other cell types during *M. tuberculosis* infection *in vivo*^12,40,74^. Indeed, NETs themselves can stimulate type I IFN production from plasmacytoid dendritic cells (pDCs)^38^ and macrophages^73^, which may contribute to the heightened type I IFN signaling in ATG5-deficient neutrophils *in vivo*. This regulatory role for ATG5 in neutrophils is autophagy-independent and specific for type I IFN-induction of NETosis and swarming. It is, therefore, possible that ATG5 is also required in other contexts where type I IFN signaling is induced, including other infections, autoimmunity disorders^38,67^, and inflammatory diseases^17,68^. In particular, NETosis has been associated with poor disease outcomes during infection with SARS-CoV-2^15^, *M. tuberculosis*^12,13^, *Pseudomonas aeruginosa*^36^ and *Francisella tularensis*^69^, which are all infections where type I IFN signaling is induced. Together our data identify ATG5 as a master regulator of downstream effects of type I IFN signaling in neutrophils, revealing a new node that may be manipulated to better control interferon-mediated disease outcomes.

## METHODS

### Mice

The *Atg5*^*fl/fl*^, *Becn1*^*fl/fl*^ and *Atg16l1*^*fl/fl*^ mice used in this study have been described previously^75–78^ and colonies are maintained in an enhanced barrier facility. LysM-Cre (#004781) and Mrp8-Cre (#021614) from the Jackson Laboratory were crossed to the floxed mice. *Ifnar1*^*-/-*79^ mice were a gift from Dr. Ashley Steed at Washington University School of Medicine and *Padi4*^*-/-*^ mice (Jax #030315) were purchased from Jackson Laboratory and bred to *Atg5*^*fl/fl*^*-LysM-Cre* mice. Male and female littermates (aged 6-12 weeks) were used and were subject to randomization. The mice were housed and bred at Washington University in St. Louis in specific pathogen-free conditions in accordance with federal and university guidelines, and protocols were approved by the Animal Studies Committee of Washington University.

### Bacterial strain and culturing

*M. tuberculosis* Erdman expressing GFP (GFP-*Mtb*^33,34^) was used in all experiments except the bulk RNA-sequencing experiment when WT Erdman was used. *M. tuberculosis* was cultured at 37°C in 7H9 (broth) or 7H11 (agar) (Difco) medium supplemented with 10% oleic acid/albumin/dextrose/catalase (OADC), 0.5% glycerol, and 0.05% Tween 80 (broth). Cultures of GFP-*Mtb* were grown in the presence of 20cg/mL kanamycin to ensure plasmid retention.

### Infection of mice with *M. tuberculosis* and measurement of bacterial burden in the lungs

*M. tuberculosis* cultures in logarithmic growth phase (OD600 = 0.5–0.8) were washed with PBS + 0.05% Tween-80, sonicated to disperse clumps, and diluted in sterile water before delivering 100 CFUs of aerosolized *M. tuberculosis* per lung using an Inhalation Exposure System (Glas-Col). Within 2 hours of each infection, lungs were harvested from at least two control mice, homogenized, and plated on 7H11 agar to determine the input CFU dose. At 14 and 21 dpi, *M. tuberculosis* titers were determined by homogenizing the superior, middle, and inferior lobes of the right lung and plating serial dilutions on 7H11 agar. Colonies were counted after 3 weeks of incubation at 37°C in 5% CO2

### Bulk RNA-Seq analysis of neutrophils from the lungs of *M. tuberculosis* infected mice

Neutrophils were isolated from the lung at 14dpi by Ly6G enrichment according to the manufacturer’s instructions (Miltenyi, 130-120-337). Briefly, the lung was perfused with PBS and digested for 45 minutes with 625μg/mL collagenase D (Roche, 11088875103) and 75U/mL DNase I (Sigma, D4527). Cells were quenched with MACS buffer and passed through a 70μM filter. RBCs were lysed with ACK lysis buffer (Gibco) and quenched with MACS buffer. Neutrophils were enriched by using an anti-Ly6G positive selection kit (Miltenyi, 130-120-337). Flow cytometry was performed to assess neutrophil purity by staining the neutrophils at a 1 in 200 dilution for Ly6G (BioLegend, 127606), Gr1 (BioLegend, 108416), Ly6C (BioLegend, 128008), CD11c (BioLegend, 117323) and CD11b (BioLegend, 101228) as described below **(Fig. S4)**. RNA was extracted from neutrophils using TRIzol Reagent (Invitrogen). Samples were prepared according to library kit manufacturer’s protocol, indexed, pooled, and sequenced on an Illumina NovoSeq. Demultiplexing was performed using fastq-multx^80^ software with a maximum of one mismatch in the indexing read. RNA-seq reads were then aligned to the Ensembl release 76 primary assembly with STAR^81^ version 2.5.1a1. Gene counts were derived from the number of uniquely aligned unambiguous reads by Subread:featureCount^82^ version 1.4.6-p52. Isoform expression of known Ensembl transcripts were estimated with Salmon^83^ version 0.8.23. Sequencing performance was assessed for the total number of aligned reads, total number of uniquely aligned reads, and features detected. The ribosomal fraction, known junction saturation, and read distribution over known gene models were quantified with RSeQC version 2.6.24.

All gene counts were then imported into the R/Bioconductor package EdgeR^84^ and TMM normalization size factors were calculated to adjust for samples for differences in library size. Genes were filtered using the filterByExpr function to remove lowly expressed genes. A multidimensional scaling plot was created using the plotMDS function. Differentially expressed genes were then identified using the exactTest function in EdgeR with a Benjamini-Hochberg adjustment method to identify all significantly expressed genes with an adjusted p-value < 0.05. The rank stat of all differentially expressed genes was calculated using qnorm(1-PValue/2)*sign(logFC). GSEA was then performed using this ranked list of genes against a database of gene sets gathered from MSigDB and Reactome.

### Flow cytometry of infected lungs

Lungs were perfused with sterile PBS and digested for 45 minutes with 625μg/mL collagenase D (Roche 11088875103) and 75U/mL DNase I (Sigma D4527). Cells were quenched with PBS + 2% Heat Inactivated FBS (HI-FBS) + 2mM EDTA and passed through a 70μM filter. Cells were suspended in PBS + 2% HI-FBS + 2mM EDTA in the presence of Fc receptor blocking antibody (BioLegend, 101302, 1:400) and stained with antibodies at a 1:400 dilution against the following mouse markers: CD11b (BioLegend, 101259), CD45 (BioLegend, 103128), Ly6G (BioLegend,127627 or 127610) and Zombie-NIR (BioLegend, 423105). Cells were stained for 20 minutes at 4°C, washed and then fixed in 4% paraformaldehyde (Electron Microscope Sciences) for 20 minutes at room temperature. H3Cit staining of fixed surface labelled cells was performed by incubating the cells overnight at 4°C in PBS 2% HI-FBS + 2mM EDTA + 0.1% saponin with monoclonal anti-H3Cit antibody (Abcam, ab219407, 1:600). Cells were washed and then incubated with donkey anti-rabbit AF555 (Invitrogen, 1:200) for 1 hour at room temperature in PBS 2% HI-FBS + 2mM EDTA + 0.1% saponin. Flow cytometry was performed on an LSR-Fortessa (BD Bioscience) or Aurora (Cytek Biosciences, with 4 laser 16V-14B-10YG-8R configuration) and analyzed using FlowJo software (Tree Star). Absolute cell counts were determined using Precision count beads (BioLegend) or by volumetric-based assessment on the Aurora. Gating strategies to identify neutrophils are in **Figure S4**.

### Neutrophil isolation, infection with *M. tuberculosis* and bacterial CFU determination

Neutrophils were isolated from the bone marrow of 6-12 week old male and female mice by Percoll gradient as previously reported^85^. Briefly the bones were flushed with HBSS and 20mM HEPES (Sigma), the RBCs were lysed through osmotic lysis and the remaining cells were filtered through a 70μM filter (CellTreat), washed, resuspended in HBSS and 20mM HEPES and loaded onto a 62% (v/v) Percoll gradient (GE Healthcare #17-0891-01) in HBSS and 20mM HEPES. The neutrophils were isolated by differential centrifugation. Neutrophils were washed and resuspended in RPMI + 10% heat-inactivated at 70°C (HI70) FBS + 1mM CaCl_2_ + 1mM MgCl_2_ and counted using an automated cell counter. Viability was assessed through Trypan exclusion. Cell count was determined using a Countess 3 (Invitrogen). Neutrophils were seeded onto 12mm glass coverslips (Fisher Scientific, #1 thickness) in 24-well plates (Corning) and either treated with 5μM ionomycin, normal mouse serum for mock treatment or infected with serum opsonized *M. tuberculosis* at an MOI of 10. GFP-*Mtb* in logarithmic growth phase (OD600 = 0.5–0.8) was washed with HBSS and 20mM HEPES, sonicated to disperse clumps, pelleted at 55xg for 10 minutes to remove clumps and opsonized with 5% normal mouse serum at room temperature for 30 minutes prior to infecting neutrophils. Cells were spun at 200xg for 10 minutes and then incubated at 37°C 5% CO_2_ for 4-18 hours. After incubation coverslips were removed from the media and placed in 4% PFA for 20 minutes for fixation prior to staining. To enumerate bacterial burden at 4 and 18 hpi Triton x-100 (Sigma, final concentration 0.05%) was added to *M. tuberculosis* infected neutrophils without removing media. Lysates were serially 10-fold diluted in 0.05% Tween 80 in PBS, plated onto 7H11 agar (Difco, supplemented with 10% OADC, 0.5% glycerol) plates and incubated at 37°C in 5% CO_2_ for 21 days until colonies were counted.

### Immunofluorescence microscopy of *in vitro* neutrophil cultures

The coverslips were washed with PBS to remove the PFA, permeabilized with 0.1% Saponin (Sigma) and blocked in 2% heat-inactivated-FBS for 1 hour at room temperature followed by incubation with primary antibodies polyclonal rabbit anti-H3Cit (Abcam, ab5103, 1μg/mL) and goat anti-Myeloperoxidase (R&D systems, AF3667, 1:500) at 4°C for 5 hours. Cells were washed with PBS and incubated with donkey anti-rabbit IgG AF555 (Invitrogen, A32794, 1:200) and donkey anti-goat IgG AF647 (Invitrogen, A32849, 1:200) antibodies and 5μg/mL Hoechst 33342 (Molecular Probes) for 1 hour at room temperature. Cells were washed with PBS and then mounted using ProLong Diamond Antifade Mountant (Thermo Fisher). Cells were imaged using a Nikon A1Rsi Confocal Microscope with NIS software at the WUCCI. Images were quantified using Fiji Image J software.

### Scanning electron microscopy of lung tissue

Mice were anesthetized and lung tissue was fixed for electron microscopy by perfusing the lungs of *M. tuberculosis* infected mice with warm Ringer’s solution containing xylocaine (0.2mg/mL) and heparin (Sigma, 20 units/mL) followed by warm 0.15M cacodylate buffer pH 7.4 containing 2.5% glutaraldehyde, 2% formaldehyde with 2mM calcium chloride. The lungs were removed and placed in 0.15M cacodylate buffer pH 7.4 containing 2.5% glutaraldehyde, 2% formaldehyde with 2mM calcium chloride overnight at 4°C. Post fixation, tissue samples were rinsed in 0.15M cacodylate buffer 3 times for 10 minutes each and subjected to a secondary fixation in 2% osmium tetroxide/1.5% potassium ferrocyanide in 0.15M cacodylate buffer for one hour. Samples were then rinsed in ultrapure water 3 times for 10 minutes each and stained in an aqueous solution of 1% thiocarbohydrazide for 20 minutes. After rinsing the samples in ultrapure water 3 times for 10 minutes each, the tissue was once again stained in aqueous 2% osmium tetroxide for one hour, rinsed in ultrapure water 3 times for 10 minutes each, and stained overnight in 1% uranyl acetate in the refrigerator at 4°C. The samples were then rinsed in ultrapure water 3 times for 10 minutes each and *en bloc* stained for 1 hour with 20 mM lead aspartate. After staining was complete, samples were briefly washed in ultrapure water, dehydrated in a graded ethanol series (50%, 70%, 90%, 100% x3) for 10 minutes in each step, and then fractured under liquid nitrogen. Fractured pieces of tissue were loaded into a critical point drier (Leica EM CPD 300, Vienna, Austria) which was set to perform 12 CO_2_ exchanges at the slowest speed. Once dried, samples were mounted with fractured surface facing up on aluminum stubs with carbon adhesive tabs and coated with 10 nm of carbon and 12 nm of iridium (Leica ACE 600, Vienna, Austria). SEM images were acquired on a FE-SEM (Zeiss Merlin, Oberkochen, Germany) at 1.5 kV and 0.1 nA.

### Scanning electron microscopy of neutrophils

Neutrophils on coverslips were incubated in 0.15M cacodylate buffer pH 7.4 containing 2.5% glutaraldehyde, 2% formaldehyde with 2mM calcium chloride overnight at room temperature. Post fixation, coverslips were rinsed in 0.15 M cacodylate buffer 3 times for 10 minutes each followed by a secondary fixation in 1% osmium tetroxide in 0.15 M cacodylate buffer for 1 hour in the dark. The samples were then rinsed 3 times in ultrapure water for 10 minutes each and dehydrated in a graded ethanol series (10%, 30%, 50%, 70%, 90%, 100% x3) for 10 minutes each step. Once dehydrated, the coverslips were loaded into a critical point drier (Leica EM CPD 300, Vienna, Austria) set to perform 12 CO_2_ exchanges at the slowest speed. Once dried, coverslips were mounted on aluminum stubs with carbon adhesive tabs and coated with 10 nm of carbon and 6 nm of iridium (Leica ACE 600, Vienna, Austria). SEM images were acquired on a FE-SEM (Zeiss Merlin, Oberkochen, Germany) at 1.5 kV and 0.1 nA.

### Immunofluorescence histology of mouse lung tissue

The post-caval lobe of the lung was inflated with 4% paraformaldehyde (PFA) and submerged in 4% PFA overnight at room temperature to inactivate *M. tuberculosis*. The lung tissue was then washed twice with PBS to remove the PFA. The lung tissue was incubated with PBS overnight at 4°C, transferred to increasing concentrations of saccharose (5-20%) and embedded in 20% saccharose/Tissue-Tek O.C.T (2:1). 5μm cryo-sections were prepared with a Leica cryostat at -19°C. Tissue sections were hydrated with PBS followed by 5 minutes of exposure to 1% SDS. The tissue was washed with PBS followed by blocking with 2% BSA and 2% HI-donkey serum in PBS for 1 hour at room temperature. Next the tissue was incubated with rabbit monoclonal anti-H3Cit antibody (Abcam, ab219407, 1:600) overnight at 4°C in a humidified chamber. The sections were washed with PBS followed by incubation with conjugated AF555 goat anti-rabbit (Invitrogen, 1:200) and AF647-Ly6G (BioLegend, 1A8, 1:200) antibodies and 5 μg/mL Hoechst 33342 (Molecular Probes) for 2 hours in a humidified chamber protected from light at room temperature. Sections were washed with PBS, autofluorescence was quenched using Vector TrueVIEW (Vector, SP-8400-15) and sections were mounted using diamond-antifade mountant (Thermo Fisher). Consecutive lung sections were stained as above or lacking the monoclonal anti-H3Cit antibody to monitor signal specificity **(Fig. S4)**. Z-stack images of regions of high cellular infiltrate were acquired on a Nikon A1Rsi confocal microscope coupled with NIS software under 20x or 60x objective. Z-stack images under 10x objective were taken using the Nikon A1Rsi confocal microscope coupled with NIS software creating a mosaic with 10% overlap to cover the entire lung section. Confocal images were quantified using Fiji particle detection where after fusing the Z-stacks, 8-bit images were assessed for total area (H3Cit and Ly6G), percent area (H3Cit and Ly6G) and average particle size (Ly6G) at the following threshold limits: 65 for H3Cit and 65 for Ly6G.

### NET quantification

Fiji software was used to quantify the number of H3Cit positive neutrophils using the threshold settings of 120 for DAPI and 100 for H3Cit for images taken with a 60x objective lens on a Nikon A1Rsi Confocal Microscope. H3Cit^+^ NETs were quantified using a Ridge Detection plugin of Fiji software with a threshold of 100 for H3Cit of images taken with a 60x objective lens on a Nikon A1Rsi Confocal Microscope. The sizes of neutrophil aggregates were measured using a Ridge Detection plugin of Fiji software with a threshold setting of 120 for the Hoechst channel of confocal Nikon A1Rsi images taken with a 20x objective lens.

### Chemical inhibition of LTB4 signaling

LTB4 signaling was inhibited by treating bone marrow neutrophils with 5-lipoxygenase inhibitor Zileuton^52^ (25μM, Cayman Chemical) or receptor antagonist U75302^52^ (10μM, Cayman Chemical) where 4e5 neutrophils (in RPMI with 10% HI70 FBS, 1mM CaCl_2_ and 1mM MgCl_2_) seeded onto 12mm glass coverslips in 24 well plates (Falcon) were pretreated for 30 minutes prior to GFP-*Mtb* infection. Neutrophils were treated with vehicle (0.036% DMSO), 25μM Zileuton or 10μM U75302 throughout the 4 hours of infection. At 4 hpi coverslips were removed from media, fixed in 4% PFA for 20 minutes and processed for confocal microscopy as above.

### ELISAs

The media from neutrophils in a 24-well plate seeded on glass coverslips at 4e5 cells/coverslip mock treated or infected with *M. tuberculosis* was used to perform a LTB4 ELISA (Cayman Chemical) to quantify the amount of LTB4 or a CXCL2 ELISA (R&D Systems) to quantify the amount of CXCL2 produced by neutrophils during *M. tuberculosis* infection or mock treatment at 4hpi. Spent media was filtered through a 0.22μM filter to remove *M. tuberculosis*. The ELISAs were performed according to manufacturer’s instructions.

### Western blot

Bone marrow neutrophils were isolated as stated above and one million neutrophils were seeded per well in a 24 well plate. Neutrophils were mock treated with NMS or infected with GFP-*Mtb* as stated above. At 4 hpi media was removed from the well by pipetting and the neutrophils were lysed in 30uL of 1X RIPA buffer (Cell Signaling Technology, 9806) containing cOmplete EDTA-free protease inhibitor cocktail (Sigma, 11873580001), 1mM PMSF (GoldBio) and phosphatase inhibitor cocktails (Sigma, P5726, P0044) and immediately boiled for 20 minutes. Protein samples were diluted with 4x Laemmli buffer, resolved on a 4-12% NuPAGE Bis-Tris gel (Thermo Fisher Scientific), transferred to nitrocellulose (Bio-Rad, 1620097) and detected with the following antibodies anti-phospho-IRF3 (Cell Signaling Technology, 4D4G), anti-phospho-TBK1 (Cell Signaling Technology, D52C2), anti-Actin (Cell Signaling Technology, 4970S) and goat-anti-rabbit-horseradish peroxidase (HRP) (Thermo Fisher Scientific, 31460). HRP was detected with ECL Prime (GE Healthcare, RPN2232). The blot was first probed for pIRF3, stripped and probed for phospho-TBK1, stripped and probed for ACTIN. The blot was stripped by exposure to 62.4mM Tris-HCl pH 6.8, 2% SDS and 0.7% β-mercaptoethanol for 30 minutes at 50°C and washing 4 times with 1X TBS 0.05% Tween-20 for 5 minutes each. The blot was then blocked in 2% BSA in 1X TBS and probed for pTBK1.

### Data and statistics

All experiments were performed at least twice using a minimum of three biological replicates. When shown, multiple samples represent biological (not technical) replicates of mice randomly sorted into each experimental group. No blinding was performed during animal experiments. Animals were only excluded when pathology unrelated to *M. tuberculosis* infection was present (i.e. bad teeth). Determination of statistical differences was performed with Prism (GraphPad Software, Inc.) using log-rank Mantel-Cox test (survival), unpaired two-tailed t-test (two groups with similar variances), or one-way ANOVA with Šídák Multiple Comparison test (to compare more than two groups). When used, center values and error bars represent the mean +/- SD. In all figures, all significant differences are indicated by asterisks: * *P<0*.*05, ** P<0*.*01, *** P<0*.*001, **** P<0*.*0001*. Non-significant comparisons of particular interest are noted as ns (not significant).

## Supporting information

Supplemental Information Combined

## ACKNOWLEDGEMENTS

This work was supported by NIH grants R01 AI132697 and U19 AI142784, a Burroughs Wellcome Fund Investigators in the Pathogenesis of Infectious Disease Award, and the Philip and Sima Needleman Center for Autophagy Therapeutics and Research to C.L.S., a Potts Memorial Foundation postdoctoral fellowship to R.L.K., a National Science Foundation Graduate Research Fellowship DGE-1143954 and the NIGMS Cell and Molecular Biology Training Grant GM007067 to J.M.K. We acknowledge the assistance of Dr. Sanja Sviben and Dr. James Fitzpatrick at the Washington University Center for Cellular Imaging (WUCCI) in electron microscopy studies, which is supported by Washington University School of Medicine, The Children’s Discovery Institute of Washington University and St. Louis Children’s Hospital (CDI-CORE-2015-505 and CDI-CORE-2019-813) and the Foundation for Barnes-Jewish Hospital (3770). We also thank the Genome Technology Access Center at the McDonnell Genome Institute at Washington University School of Medicine for RNA sequencing and analysis. The Center is partially supported by NCI Cancer Center Support Grant #P30 CA91842 to the Siteman Cancer Center and by ICTS/CTSA Grant# UL1TR002345 from the National Center for Research Resources (NCRR), a component of the National Institutes of Health (NIH), and NIH Roadmap for Medical Research. This publication is solely the responsibility of the authors and does not necessarily represent the official view of NCRR or NIH. The model in Supplementary Figure 3E was created with BioRender.com.

## AUTHOR CONTRIBUTIONS

R.L.K. and C.L.S. wrote the manuscript. Experiments were designed and performed by R.L.K., C.S.C., A.S., and J.M.K. under the guidance of C.L.S. RNA sequencing data was analyzed by Y.M., E.E. and A.N.M. Mice were bred and maintained by D.K. Funding and direction were provided by C.L.S.

