## Supplemental Information Combined for "ATG5 suppresses type I IFN-dependent neutrophil swarming and NETosis"

#### Supplemental Table and Figure Legends

**Supplemental Table 1. Differentially expressed genes in neutrophils isolated from the lungs of *Atg5*$^{fl/fl}$*-LysM-Cre* and *Atg5*$^{fl/fl}$ mice at 14 dpi.** Differentially expressed genes comparing *Atg5*$^{fl/fl}$*-LysM-Cre* to *Atg5*$^{fl/fl}$ were identified using the exactTest function in EdgeR with a Benjamini-Hochberg adjustment with an adjusted p-value < 0.05. We note that the increased reads for the *Atg5* transcript in neutrophils from *Atg5*$^{fl/fl}$*-LysM-Cre* mice map primarily to exon 7, a region outside the area flanked by the loxP sites.

### Supplemental Figure 1

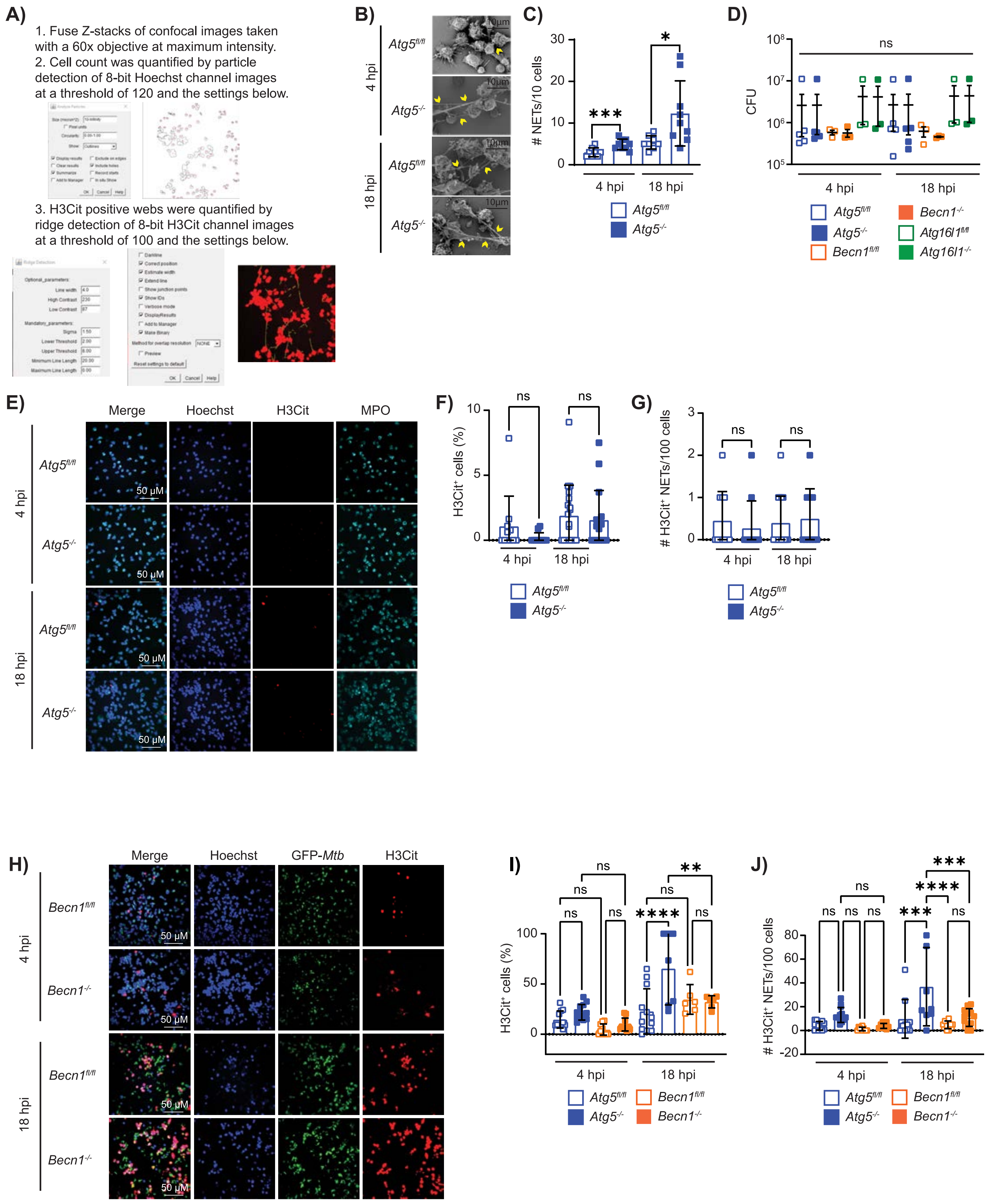

**Supplemental Figure 1. *In vitro* experiments with neutrophils isolated from the bone marrow of *Atg5$^{fl/fl}$*, *Atg5$^{fl/fl}$-LysM-Cre, Becn1$^{fl/fl}$*, and *Becn1$^{fl/fl}$-LysM-Cre* mice. (A)** The parameters used to measure the frequency of Hoechst$^+$ cells that were H3Cit$^+$ by Fiji particle analysis and the number of extracellular H3Cit$^+$ NETs using Fiji Ridge Detection. **(B)** Representative SEM images of mouse neutrophils infected with GFP-*Mtb in vitro* at 3000x magnification. Yellow arrows indicate NETs being released from cells. **(C)** The number of NETs released per 10 cells quantified from SEM images at 1000X magnification of neutrophils infected with GFP-*Mtb* for 4 or 18 hpi. Each datapoint represents one field with 3 fields containing 10-40 cells/field compiled from 3 independent experiments. **(D)** *M. tuberculosis* CFU recovered from cultures of neutrophils infected with GFP-*Mtb in vitro* for 4 or 18 hpi. Each datapoint is from an independent well of infected cells compiled from at least 2 independent experiments. **(E)** Representative confocal immunofluorescence microscopy images of neutrophils mock infected *in vitro* for 4 or 18 hours and stained for H3Cit, the neutrophil marker myeloperoxidase (MPO), and DNA (Hoechst) (60X magnification). **(F)** The proportion of Hoechst$^+$ cells that were H3Cit$^+$ and **(G)** the number of extracellular H3Cit$^+$ NETs released per 100 cells after mock-infection of neutrophils for 4 or 18 hours, as quantified from fluorescence microscopy experiments. **(H)** Representative confocal immunofluorescence images at 60X magnification of neutrophils infected with GFP-*Mtb in vitro* for 4 or 18 hpi and stained for DNA (Hoechst), H3Cit, and MPO. GFP-*Mtb* is also shown. **(I)** The proportion of Hoechst$^+$ cells that were H3Cit$^+$ and **(J)** the number of H3Cit$^+$ NETs released per 100 cells neutrophils at 4 and 18 hpi with GFP-*Mtb*, as quantified from fluorescence microscopy experiments. **(F-G, I-J)** Each datapoint represents data from one field, with a minimum of 6 fields containing 40-150 cells/field each compiled from at least 2 independent experiments. All graphs report the mean ± SD. Statistical differences were determined by one-way ANOVA and Šídák multiple comparison test **(D)** or student t-test **(C, F-G, I-J,** only comparing within a single timepoint). * $P < 0.05$, ** $P < 0.01$, and **** $P < 0.0001$. Differences that are not statistically significant are designated as ns.

### Supplemental Figure 2

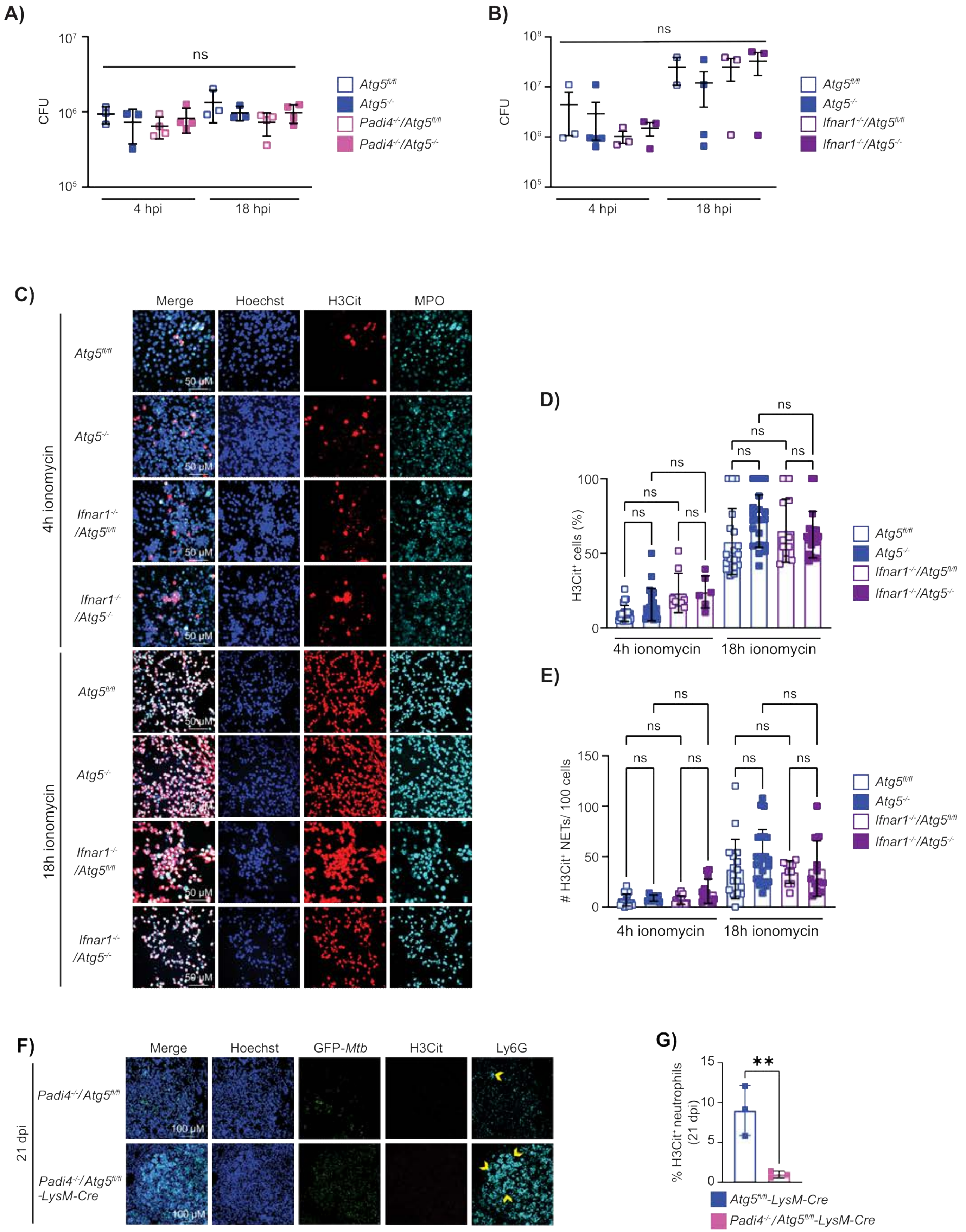

**Supplemental Figure 2. ATG5 specifically suppresses type I IFN-induced PAD4-mediated histone citrullination *in vitro* and *in vivo*. (A-B)** *M. tuberculosis* CFU recovered from cultures of mouse bone marrow neutrophils infected with GFP-*Mtb in vitro* for 4 and 18 hpi. Each datapoint is from a single well compiled from at least 2 independent experiments. **(C)** Representative confocal immunofluorescence microscopy images at 60X magnification of neutrophils treated with ionomycin for 4 or 18 hours and stained for DNA (Hoechst), H3Cit, and neutrophil marker MPO. **(D)** The proportion of Hoechst$^{+}$ cells that are H3Cit$^{+}$ and **(E)** the number of H3Cit$^{+}$ NETs released per 100 cells from neutrophils treated with ionomycin for 4 or 18 hours, based on fluorescence microscopy experiments. **(F)** Representative confocal immunofluorescence images at 20X magnification of lung sections from GFP-*Mtb* infected mice at 21 dpi probed with antibodies to detect citrullinated histone H3 (H3Cit; red), neutrophil marker Ly6G (cyan), and DNA (Hoechst; blue). GFP-*Mtb* is also shown. Yellow arrows indicate Ly6G$^{+}$ aggregates that are greater than 70 pixels in size. **(G)** The proportion of neutrophils (CD45$^{+}$Ly6G$^{+}$CD11b$^{+}$) that were H3Cit$^{+}$ at 21 dpi in the lungs of mice quantified by flow cytometry. Each data point is from an individual mouse compiled from at least two separate infection experiments. All graphs report the mean ± SD. Statistical differences were determined by one-way ANOVA and Šídák multiple comparison test where ** $P < 0.01$. Differences that are not statistically significant are designated as ns.

#### Supplemental Figure 3

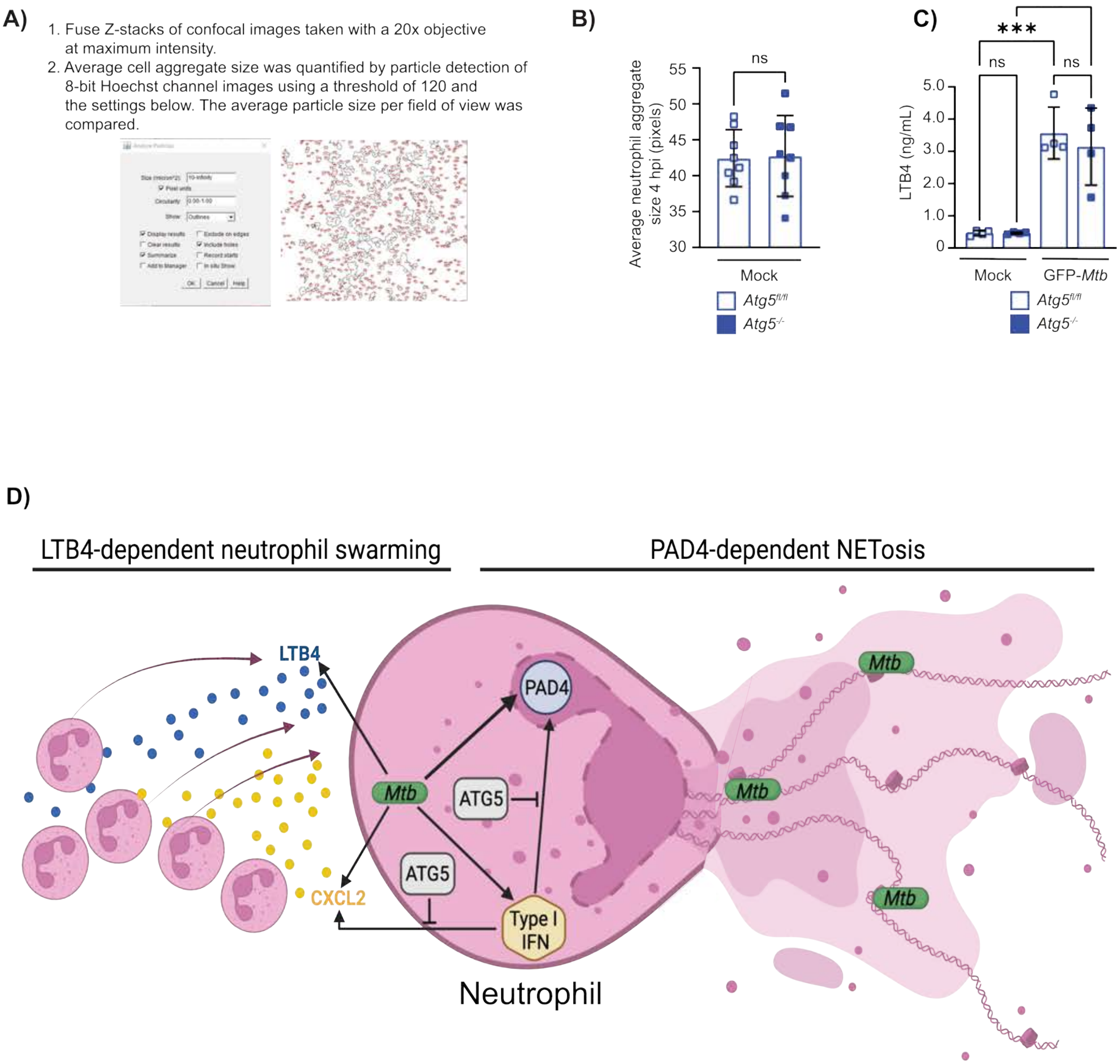


**Supplemental Figure 3. Neutrophil aggregation and LTB4 production during mock and *M. tuberculosis* infection *in vitro*, and model of ATG5 activity in neutrophils. (A)** The Fiji particle analysis parameters used to measure the average size of neutrophil aggregates. **(B)** The average neutrophil aggregate size at 4 hpi of mock treated mouse neutrophils based on fluorescence microscopy. Each datapoint represents the average neutrophil aggregate size per field, with a minimum of 8 fields containing 100-250 cells/field each compiled from at least 2 independent experiments. **(C)** The amount of LTB4 present in neutrophil culture supernatants from mock treated or GFP-*Mtb* infected mouse neutrophils at 4 hpi quantified by ELISA. Each datapoint is from an independent well of infected cells compiled from at least 2 independent experiments. Bar graphs report the mean ± SD. Statistical differences were determined by student t-test **(B)** or one-way ANOVA and Šídák multiple comparison test **(C)**. *** $P < 0.001$. Differences that are not statistically significant are designated as ns. **(D)** Model depicting how ATG5 functions in neutrophils to regulate type I IFN-induced NETosis and swarming. Created with BioRender.com.

#### Supplemental Figure 4

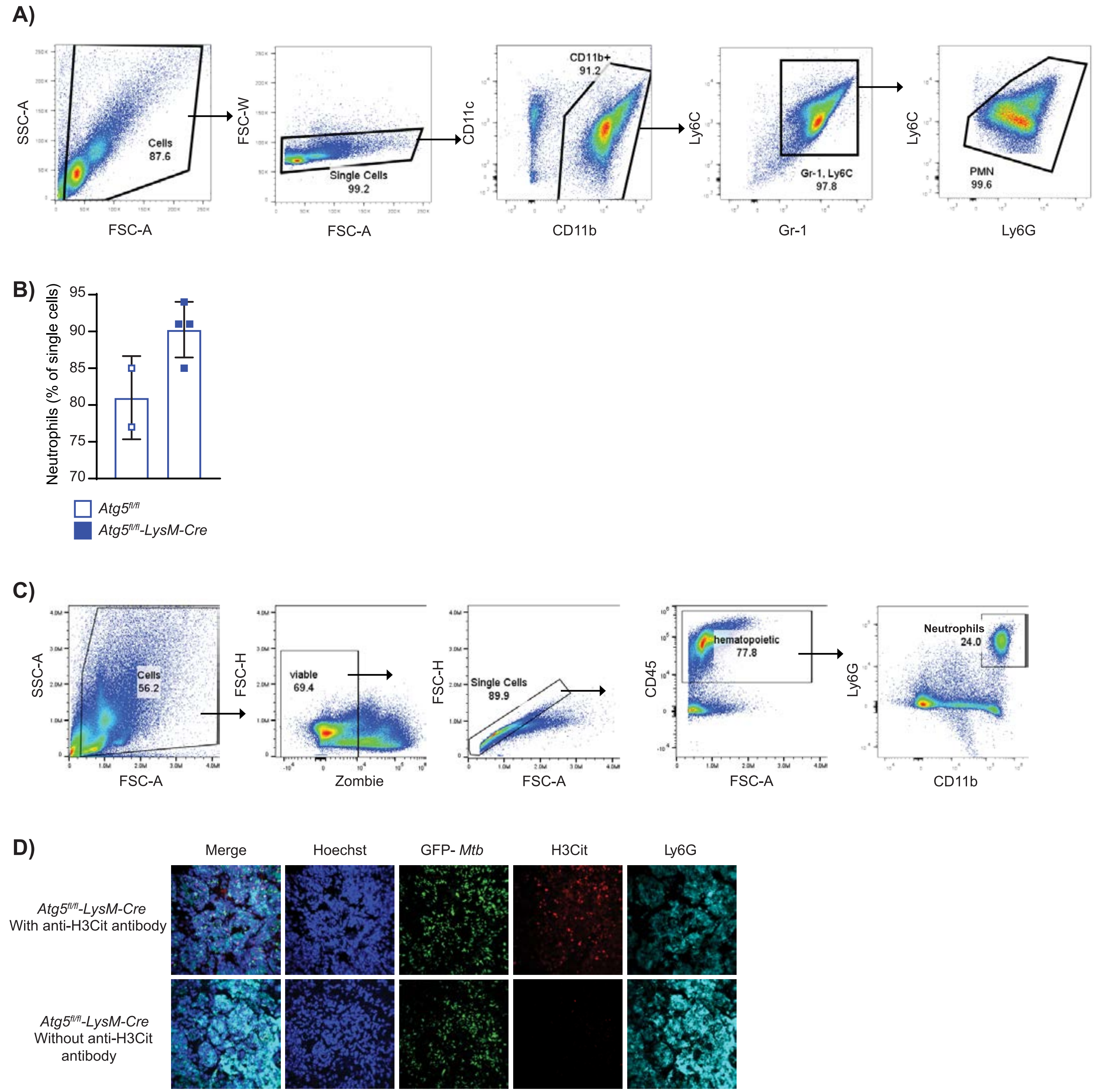


**Supplemental Figure 4. Flow cytometry assessment of neutrophil purity following anti-Ly6G selection and flow cytometry gating strategy to identify neutrophils in the lung. (A)** The gating strategy used to identify neutrophils (GR1$^{+}$Ly6G$^{+}$CD11b$^{+}$) after anti-Ly6G enrichment from the lung at 14 dpi. **(B)** The proportion of single cells that are neutrophils from *Atg5$^{fl/fl}$* and *Atg5$^{fl/fl}$-LysM-Cre* mice after anti-Ly6G enrichment from the lung at 14 dpi. Bar graphs report the mean ± SD where each data point is from an individual mouse. **(C)** The gating strategy used to identify live neutrophils (Zombie-NIR$^-$ CD45$^+$ Ly6G$^+$ CD11b$^+$) in the lung at 14 or 21 dpi. **(D)** Representative confocal immunofluorescence microscopy images of lung sections from GFP-*Mtb* infected *Atg5$^{fl/fl}$-LysM-Cre* mice at 21 dpi probed with antibodies to detect citrullinated histone H3 (H3Cit; red), Ly6G (neutrophil marker; cyan), and DNA (Hoechst; blue). GFP-*Mtb* is also shown. A consecutive section was stained with Hoechst and probed with anti-Ly6G, and anti-rabbit IgG-AF555 antibodies but was not probed with the primary anti-H3Cit antibody.
